# Ammonium Pretreatment and Addition Improve Stability of Environmental Parameters and Biogas Production under Anaerobic Co-digestion of Rice Straw and Dairy Manure

**DOI:** 10.1101/335588

**Authors:** Yun Tang, Shuzhen Zou, Di Kang

**Author notes:** Corresponding author Address: College of Life Science, China West Normal University, Nanchong 637000, Sichuan, China.

## Abstract

This paper optimized the anaerobic digestion (AD) pretreatment process, identified the relationship between stability of environmental factors and biogas production under ammonium hydroxide (NH_3_·H_2_O) pretreatment and analyzed the reason of NH_3_·H_2_O pretreatment to increase biogas production. Variable coefficients (CVs) of environmental factors were calculated to study the stability of environmental factors during AD process. The effect of initial AD environment factors on the stability of environmental factors during AD process was analyzed by redundancy analysis. Path analysis was used to analyze the response relationship the stability of environmental factors between and total biogas production (TBP). Results showed that pretreatment at 8% for 4 days, the TBP produced the highest value (302.5mL/g TS) and significantly higher than the other values (P < 0.01). NH_3_·H_2_O pretreatment had effect on the initial AD environment factors and the environment factors during AD process. Under the NH_3_·H_2_O pretreatment conditions, the stability of environment factors during AD process was affected by initial AD environment factors, while they had direct and indirect influences on the TBP. This research concluded that NH_3_·H_2_O pretreatment improved TBP via changing the initial environment of AD and the stability of environment factors during AD process, as well as the response relationship among initial AD environment factors and the stability of environment factors during AD process and biogas production, the changes improved the stability of environmental factors and made the environment more suitable for AD.

## 1 Introduction

Agricultural pollution and fossil energy crisis are now becoming two serious issues all over the world [1]. Anaerobic digestion (AD) usually refers to the microbial conversion of organic material to biogas, which mainly consists of methane and carbon dioxide, as a carbon-neutral alternative to fossil fuels whilst destroying pathogens and removing odors [2]. Thus, it has been used to alleviate the issues of environment pollution and energy crisis [3]. However, there are two factors limited AD. First, the wax layer on the epidermis of straw slows the absorption of water and limits the disintegration of lignocelluloses. Second, the high C/N and lignin in straw made AD has a hard starting. Results have showed that pretreatment method and co-AD could make up the shortfall of AD and enhance the methane potential and/or AD rate, improving digester performance [4-6]. China had the most people in the world, but living scattered and the way of land use is family contract, so the state and the farmers see rural household biogas development as an important approach to help solve environmental and rural energy problems. Pretreatments used in rural household biogas must be simple, time-saving, inexpensive and environmental friendly. NH_3_·H_2_O pretreatment are carried out under milder conditions, some of them even at ambient temperature [7]. Thus, the NH_3_·H_2_O pretreatment may be more suitable for rural household biogas in China.

AD process model indicated that there were three stages in AD process, which were liquid stage, acidification stage, biogas stage [8]. In the first stage, the complex organic matter was decomposed into dissolved organic matter. In the second stage, the dissolved organic matte was metabolized to monosaccharide, then it was decomposed to volatile fatty acid (VFA). The VFA was dissolved by methane bacteria to produce biogas [9]. Thus, the total organic carbon (TOC), reducing sugar and VFA were the keys carbon sources in the three stages in AD process. The growth rate of microorganisms was significantly affected by pH changing [10], because a significant positive correlation (P<0.01) was found between hydrolysis rate and pH [11], Yin et al. [12] also showed that pH displayed the maximum comprehensive influence with total biogas production (TBP) in AD. Cellulase activity can ensure the optimal growth and activity of various types of microorganisms, therefore, biomass can be more resistant to shock loading [13]. Thus, pH and cellulase activity were the two important micro-ecological environment factors in AD process, and they may have the most effect on the first stage and third stage in AD process. Previous studies mostly focused on third stage of AD [14,15]. For examples, at the beginning of the AD, VFA content was low, and thus, VFA was produced by hydrolysis and acid bacteria, thereby increasing the VFA content and decreasing the pH, along with the digestion, VFA, other acid, and dissolved nitrogen-containing compounds were broken into CH_4_ [16], thereby decreasing the VFA content and increasing the pH [17]. Individual VFA were measured to examine process performance in AD [18], and the average daily biogas production (DBP) was inversely proportional with VFA and direct proportion with pH [19], for VFA content and pH had an effect on buffer capacity of AD and improve the AD stability [20]. However, the TOC, reducing sugar and cellulase activity that affected the first stage and second stage and their effect on TBP were few researched.

Our results have showed that initial environmental factors in AD and environmental factors during AD process affected the DBP and TBP, respectively [21]. Mao et al. [22] indicated initial environmental factors had effect on the environmental factors during AD process. And may researchers have showed that a good stability of AD process was a precondition to perform the AD and produce biogas successfully. The stability of DBP [23], VFA [24], pH [25] and microorganism [26] has been used to measure the stability of system in AD process. Generally speaking, previous results pointed out that the TBP was higher and the peak DBP was higher, we thought the system of AD was more stability, and in statistics, stability was expressed by coefficient variation (CV) [27], which indicated that the stability of AD system may affected by the CV of environmental factors in AD system, such as TOC, reducing sugar, VFA, cellulase activity and pH. However, there are three points that important but have not been studied. First, the CVs of environmental factors in AD system were calculated rarely in previous researches. Second, little previous researches used the CVs of environmental factors during AD system to analyze the relationship between the stability of AD system and TBP. Last, the relationship between initial characteristics in AD and CV of characteristics in AD process and the effect of CVs of characteristics during AD process on TBP were not unclear. Thus, further investigation is required to focus on the TOC, reducing sugar, VFA, pH, cellulase activity and their effect on the CV of environmental factors during AD process, as well as clarify the effect of CV of environmental factors during AD process on the TBP. In addition, it is necessary to discuss the reasons of NH_3_·H_2_O pretreatment to improve the TBP based on the stability of AD system.

To address these gaps, the soluble carbon substances (TOC, reducing sugar, VFA) and the ecological factors (cellulase activity and pH) were measured in initial characteristics and process characteristics of AD, their CVs were calculated. Redundancy analysis was used to analyze the effect of initial characteristics (TOC, reducing sugar, VFA, pH and cellulase activity) on the CVs of process characteristics (TOC, reducing sugar, VFA, pH, cellulase activity and DBP). The effect of CVs of process characteristics on the TBP was analyzed by path analysis. The aims of this paper included optimizing the NH_3_·H_2_O pretreatment conditions, discussing the reasons of NH_3_·H_2_O pretreatment to improve the TBP based on the factors characteristics in initial environment and process environment, as well as the stability of AD system.

## 2 Materials and Methods

### 2.1 Substrate and inoculum

Rice straw (RS) was obtained from a farm after the new harvest and smashed into lengths of 1-2 cm pieces by a 9Z-2.5 straw material shredder before the pieces were added to use. The dairy manure (DM) was gathered from a livestock farm. The inoculum was obtained from household bio-energy digesters. The chemical characterization of each substrate and the sludge tested in this study are presented in table 1. All samples were collected in triplicate, and the averages of the three measurements are presented. The farm, livestock farming and household bio-energy digesters were located in Cuixigou in Yangling, China.

**Table 1.** Characterizations of substrates used in the AD experiments

| Material | TOC<sup>a</sup> (g/kgVS) | TON<sup>a</sup> (g/kgVS) | C/N<sup>a</sup> | TS (%) | VS (%) |
| --- | --- | --- | --- | --- | --- |
| Rice straw | 45.27±0.21 | 0.62±0.10 | 73.02 | 90.00±0.18 | 83.89±0.17 |
| Dairy manure | 56.21±0.58 | 3.65±0.21 | 15.4 | 21.22±0.2 | 84.07±0.59 |
| Inoculum | 39.00±0.32 | 1.81±0.21 | 21.55 | 9.23±0.09 | 75.83±0.6 |

Note: <sup>a</sup> Dry weight basis. TOC, TON, TS and VS are the abbreviations of total organic carbon, total organic nitrogen, total solids and volatiles solids.

### 2.2 NH_3_·H_2_O pretreatment method

Putting 120mL, 160mL, 200mL and 240mL of pure water into four of 2000 mL volumetric flasks, and diluted with purified water to 2000mL. Thus, the concentrations of NH_3_·H_2_O in the volumetric flasks were 6%, 8%, 10% and 12% concentrations (V/V). Four 500g of RS samples (TS content was 91%) were weighed and putted into four of 2L beakers to further access. Then 1822g of 6%, 8%, 10% and 12% concentrations (V/V) of NH_3_·H_2_O were added into the 2L beakers, respectively, to make the TS content was 20% in the 2L beakers. Lastly, the RS with NH_3_·H_2_O was pretreated for 2 days, 4 days and 6 days, respectively, under room temperature (25°C±2°C). The pretreated RS samples were stirred once every 12 hours.

### 2.3 Anaerobic digestion method

The pretreated RS was mixed with DM to total of 56 g TS (pretreated RS and DM amounts of 28 g and 28 g), then 56 g TS and 200 g inoculum were placed in a 1 L Erlenmeyer flask, and pure water was added to a total mass of 700 g, assuring that the TS content was 8%. AD was conducted for 35 days under 35°C. Figure 1 depicts the experimental equipment. Biogas generated in figure 1 (a) was transported into the headspace of the bottle using a glass pipe (figure 1c) before entering a 15 L biogas collection device (figure 1f) through figure 1 (e). The water in figure 1 (g) was pressed out by a piston (figure 1h) and overflowed into figure 1 (j) through another glass pipe (figure 1i). The volume of the discharged water from the bottle represents the volume of the biogas generated. A 1 L measuring cylinder (figure 1j) was used to measure the water displaced from the collector each day. After 35 days of AD, the methane content was measured as shown in figure 1 (e). The headspace (figure 1c) of the reactors was flushed with pure nitrogen gas for 1 min to achieve anaerobic conditions, and the reactors (figure 1a) were immediately sealed with air impermeable rubber stoppers for AD. Three blank controls were operated to remove endogenous methane from the sludge [28].

**Figure 1.**
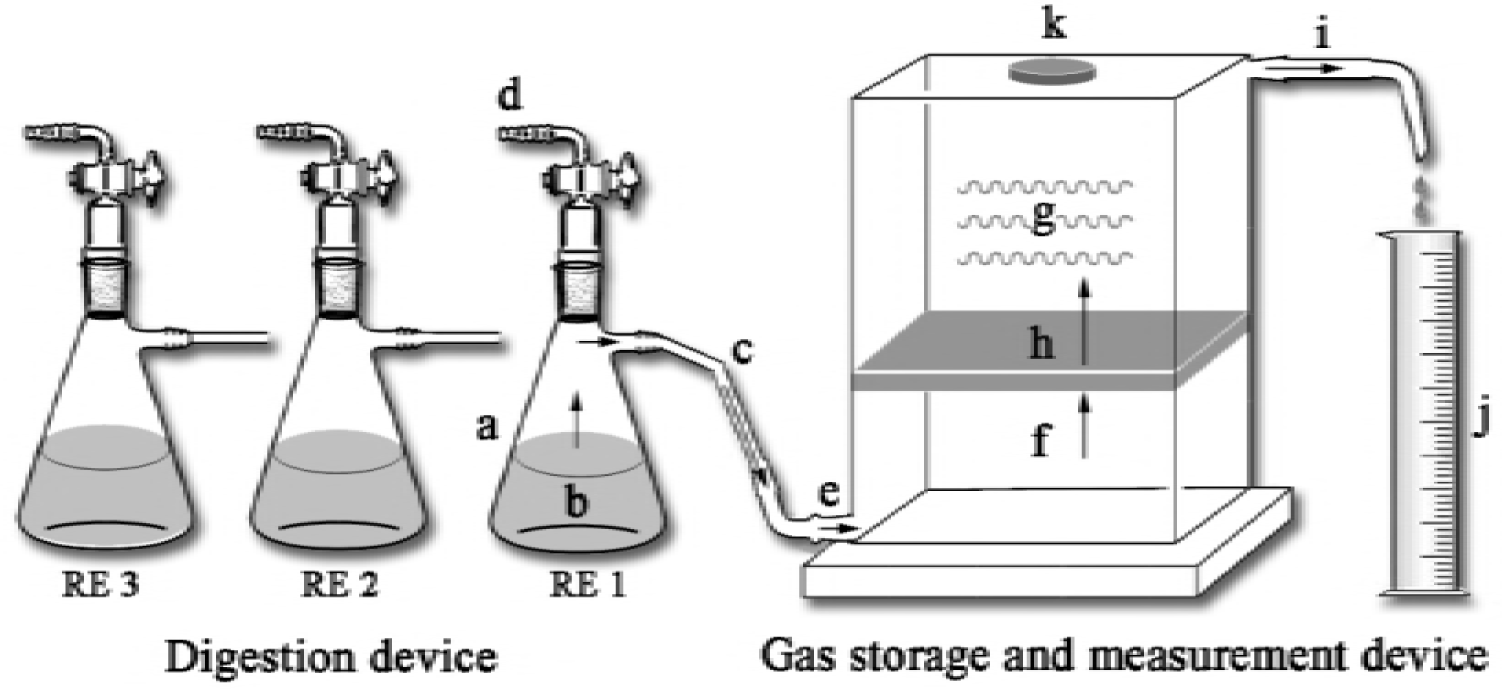
Controlled constant temperature AD device. Note: a. Digester; b. Biogas fluid and substrates; c. Airway tube; d. Taking biogas and sampling; e. Air intake; f. Biogas collecting bottle; g. Water; h. Piston; i. Aqueduct; j. Water collecting and measuring cylinder; and k. Water inlet.

**Figure 2.**
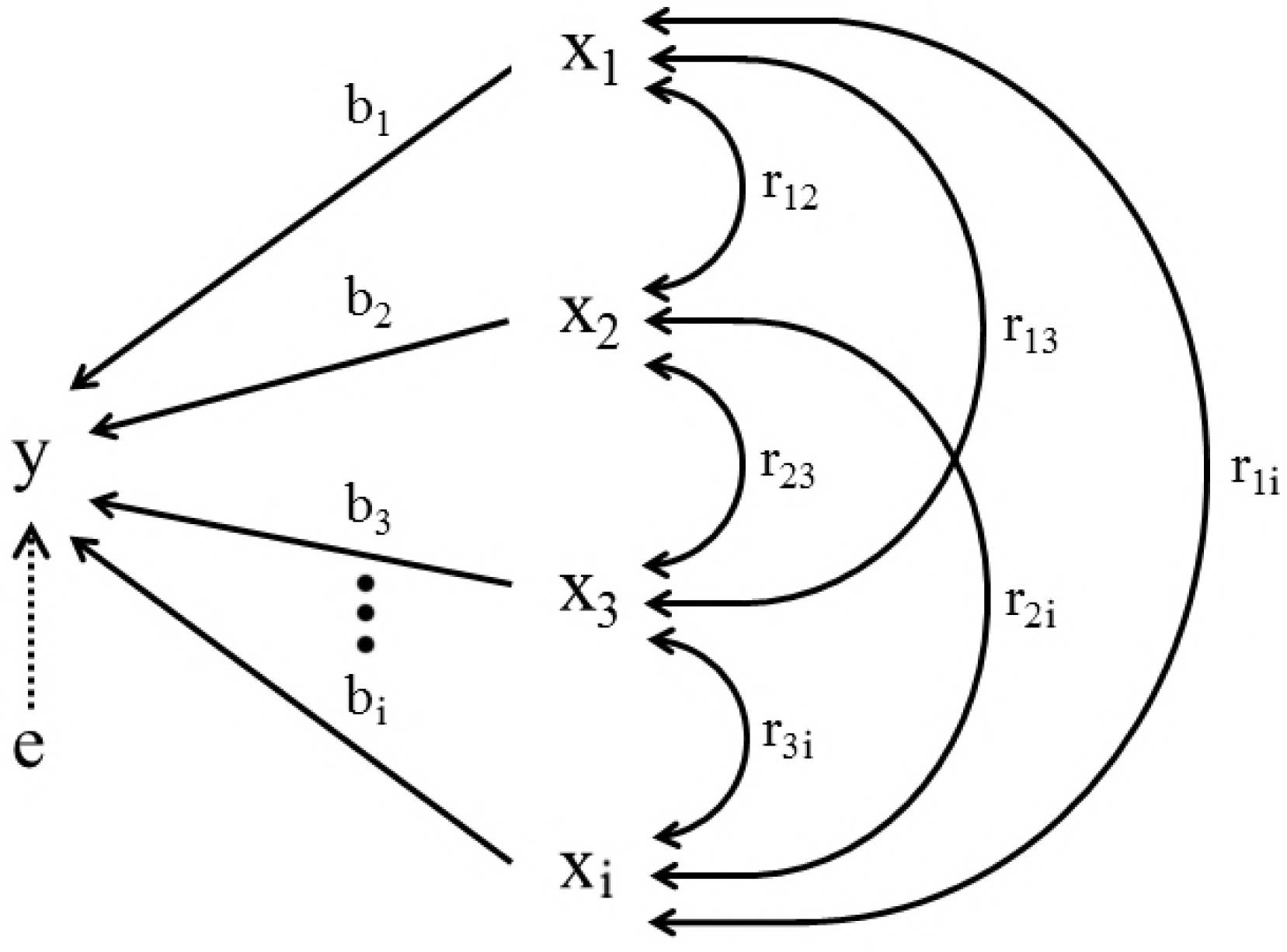
Path coefficient analysis. Note: i=7.x_1_, x_2_, x_3_, x_4,_ x_5_ and e represent the variable coefficient of total organic carbon content, reducing sugar content, volatile fatty acids content, cellulase activity, pH and other factor (s), respectively. r is correlation coefficient. b_1_, b_2_, b_3_ and b_i_ are partial regression coefficients. y represents the TBP and e is residual path coefficient.

### 2.4 Statistical analysis

#### 2.4.1 Coefficient of variable

In probability theory and statistics, the CV, also known as relative standard deviation, is a standardized measure of dispersion of a probability distribution or frequency distribution. The CV has no dimension, so it can be compared objectively. It was influenced by dispersion degree and average value of variables. The CV can prove the stability of the variables, and the greater the CV, the lower the stability. The formula for the CV is as follows:

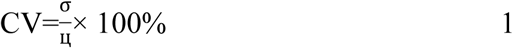

Where, CV was coefficient of variable, σand Ц was the standard deviation and mean of the population.

#### 2.4.2 Redundancy analysis

Redundancy analysis allows studying the relationship between two tables of variables Y and X. While the canonical correlation analysis is a symmetric method, Redundancy Analysis is non-symmetric. In Redundancy Analysis, the components extracted from X are such that they are as much as possible correlated with the variables of Y. Then, the components of Y are extracted so that they are as much as possible correlated with the components extracted from X. In this study, the X matrix the factors in initial AD environment, and Y matrix was the CV of environmental factors during AD process. Redundancy analysis method was used to analyze the relationship between the factors in initial AD environment and the environmental factors stability during AD process.

### 2.5 Analytical methods

The determination of total solid (TS), volatile solid and VFA content was based on methods described for routine analysis of bio-energy digestion [29]. The TOC was determined by the method described in Cuetos [30]. Total organic nitrogen was determined using the Kjeldahl Method [31]. Cellulase activity and reducing sugar content were determined by the DNS method [32], the reducing sugar content is used to measure the cellulase activity. The DBP was measured by displacement of water, and the pH was measured with by Model DLGA-1000 analyzer (Infrared Analyzer; Dafang, Beijing, China).

## 3 Results and discussion

### 3.1 The influence of NH_3_·H_2_O pretreatment on biogas production

As shown in figure 3 (a), the largest DBP for NH_3_·H_2_O pretreated samples was higher than CK (P< 0.05), and their appearance time were affected by the NH_3_·H_2_O pretreatment. It took 24 days to reach the largest DBP of 710 mL for CK. While the largest DBPs after NH_3_·H_2_O pretreatment of C_8_ d_2_, C_8_ d_4_, C_8_ d_6_, C10 d_2_, C10d_6_ and C1_2_d_6_ were 870 mL, 1330 mL, 875 mL, 930 mL, 780 mL and 780 mL reached at 20, 11, 18, 13, 15 and 14 days, respectively. We concluded that NH_3_·H_2_O pretreatment could improve the largest DBP of NH_3_·H_2_O pretreated samples and advanced their appearance time of peak DBP. May be because some of the unavailable carbon might has been converted quickly during the NH_3_·H_2_O pretreatment to soluble carbon that could be used by bacteria during AD [33], which was supported by Wei et al [34].

**Figure 3.**
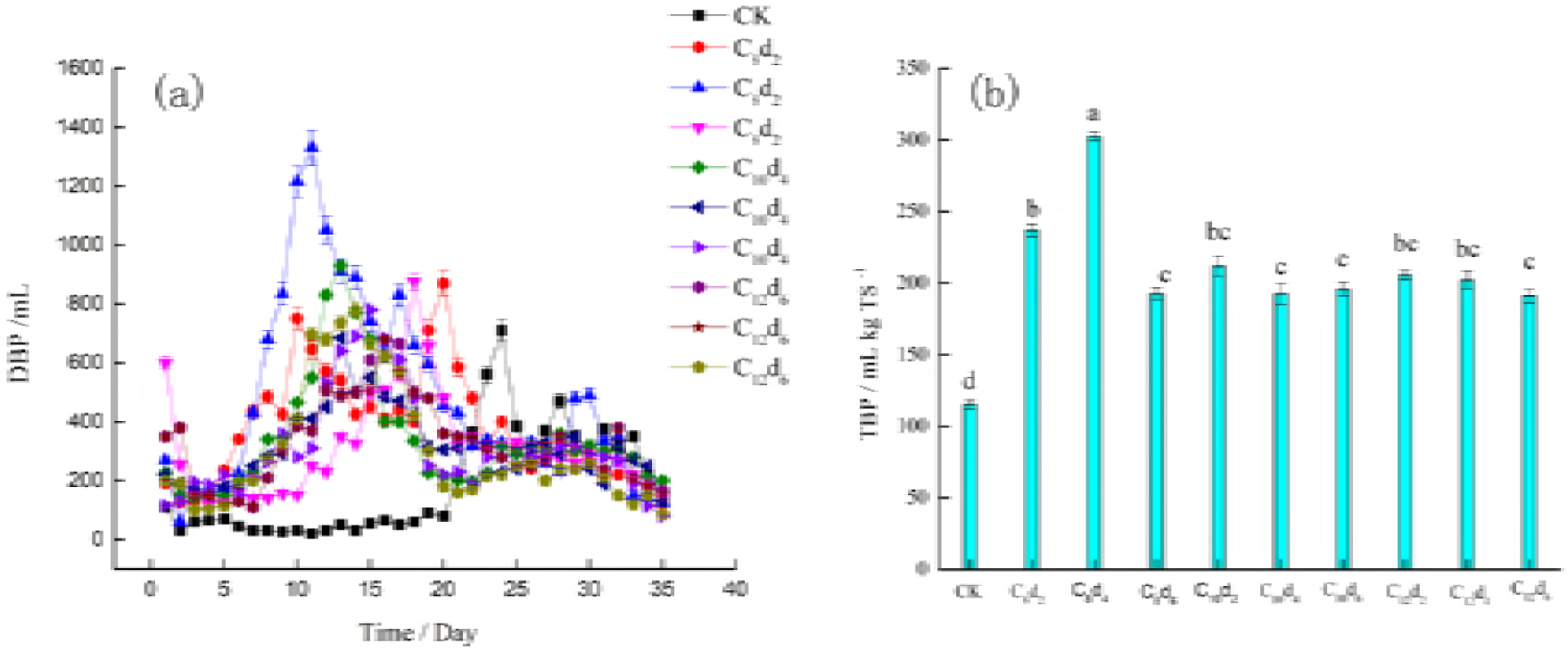
DBP and TBP of AD under NH_3_·H_2_O pretreatment. Note: CK is un-pretreated sample; C and d are the NH_3_·H_2_O pretreatment concentration and NH_3_·H_2_O pretreatment. DBP and TBP means the daily biogas production and total biogas production. The same superscripts indicate insignificant difference (P < 0.05). Different superscripts indicated significant difference (P < 0.05).

The effects of NH_3_·H_2_O pretreatment on TBP are shown in figure 3 (b). When RS was pretreated with 8% for 4 days, the TBP produced the highest value (302.5mL/g TS), which was 263% higher than that of CK (115.18 mL/gTS) and significantly higher than the other values (P < 0.01), which indicated that NH_3_·H_2_O pretreatment could improve TBP. Similarly, many previous studies have shown the same result [35,36]. For example, previous study has shown that the highest TBP of NH_3_·H_2_O pretreated straw was enhanced 1.56 times compared with that from CK. And Wei et al. [34] pointed out that the when the NH_3_·H_2_O pretreated maize straw, then it was mixed with dairy manure, the TBP significantly increased 21. 40% compared with CK (P < 0.01). Thus, we quite sure that NH_3_·H_2_O should be used for the RS pretreatment to enhance biogas production in household biogas in China, theoretically.

### 3.2 The influence of NH_3_·H_2_O pretreatment on initial environment of AD

As showed in figure 4, the contents of TOC and reducing sugar in CK were 30.23 g/L and 9.20 g/L. The contents of TOC and reducing sugar in C_12_d_2_ and C_12_d_6_ were the highest (69.46 g/L and 40.35 g/L), and significantly higher than that of CK (P<0.05). VFA content in CK was 0.712 g/L, while in the pretreated sample, the 1.092 g/L was the highest under C_8_d_4_· The result indicated that NH_3_·H_2_O pretreatment changed the initial characteristics of AD. After NH_3_·H_2_O pretreatment, the contents of TOC, reducing sugar and VFA content were higher than that of CK, because the lignin and hemicellulose was dissolved by NH_3_·H_2_O into organic matter [37], such as TOC, glucose, VFA and so on. Results have showed that the TOC content increased by approximately 478% [38] and VFA content [35], which supported our results. In CK, the pH value in initial AD of CK is 7.2, while all of treatments after NH_3_·H_2_O pretreatment, the pH were higher than that of CK, for NH_3_·H_2_O was a kind of alkaline reagents, it was left in AD system to increase the pH. After NH_3_·H_2_O, the cellulase activity of all treatments was higher than that of CK. May be because some inorganic ions had activated effect on cellulase activity in suitable concentration range and maybe NH_4_ ^+^ had an activate effect on cellulase activity [39]. Thus, we concluded that NH_3_·H_2_O pretreatment could improve the content of TOC, reducing sugar and VFA in initial characteristics of AD. As well as NH_3_·H_2_O pretreatment improved the microbial ecological environment of cellulase activity and pH.

**Figure 4.**
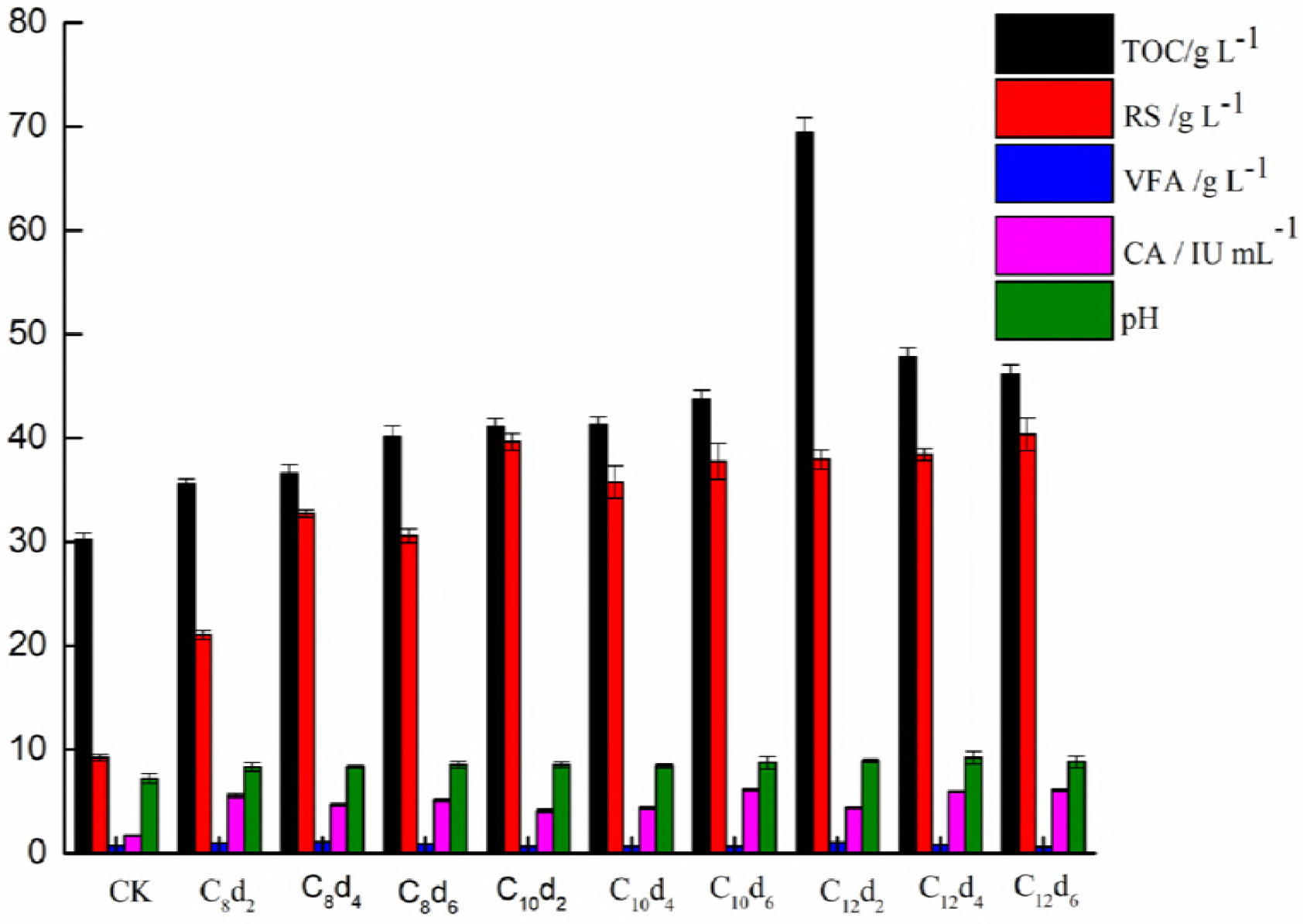
Initial environment of AD under NH_3_·H_2_O pretreatment. Note: CK is un-pretreated sample; C and d are the NH_3_·H_2_O pretreatment concentration and NH_3_·H_2_O pretreatment time. TOC and VFA mean the total organic carbon and volatile fatty acid.

### 3.3 The influence of NH_3_·H_2_O pretreatment on the environmental factors during AD process

#### 3.3.1 The influence of NH_3_·H_2_O pretreatment on the nutrient substance content during AD process

As shown in figure 5 (a), the highest TOC content of CK was 64.38 mg/L, and its built-up time appeared on the 10th day. The highest TOC contents of C_8_d_2_, C_8_d_4_, C_8_d_6_, C_10_d_2_, C_10_d_4_, C_10_d_6_, C_12_d_2_, C_12_d_4_, C_12_d_6_ were 44.77 mg/L, 37.62 mg/L, 52.50 mg/L, 49.32 mg/L, 68.77 mg/L, 58.21 mg/L, 44.54 mg/L, 60.37 mg/L and 48.35 mg/L, respectively, they appeared on days 5. It was indicated that all of the highest TOC contents of pretreated samples in AD process were lower than that of CK, but their appearance days was advanced. Because, the area of lignocellulose and cellulose content of RS increased, which made the hydrolysis stage accelerate, thus appearance peak days of TOC were advanced [40], but the some lignin and hemicellulose was decomposed by NH_3_·H_2_O pretreatment, the carbon material reduced [35], thus the TOC content in pretreated samples was lower.

**Figure 5.**
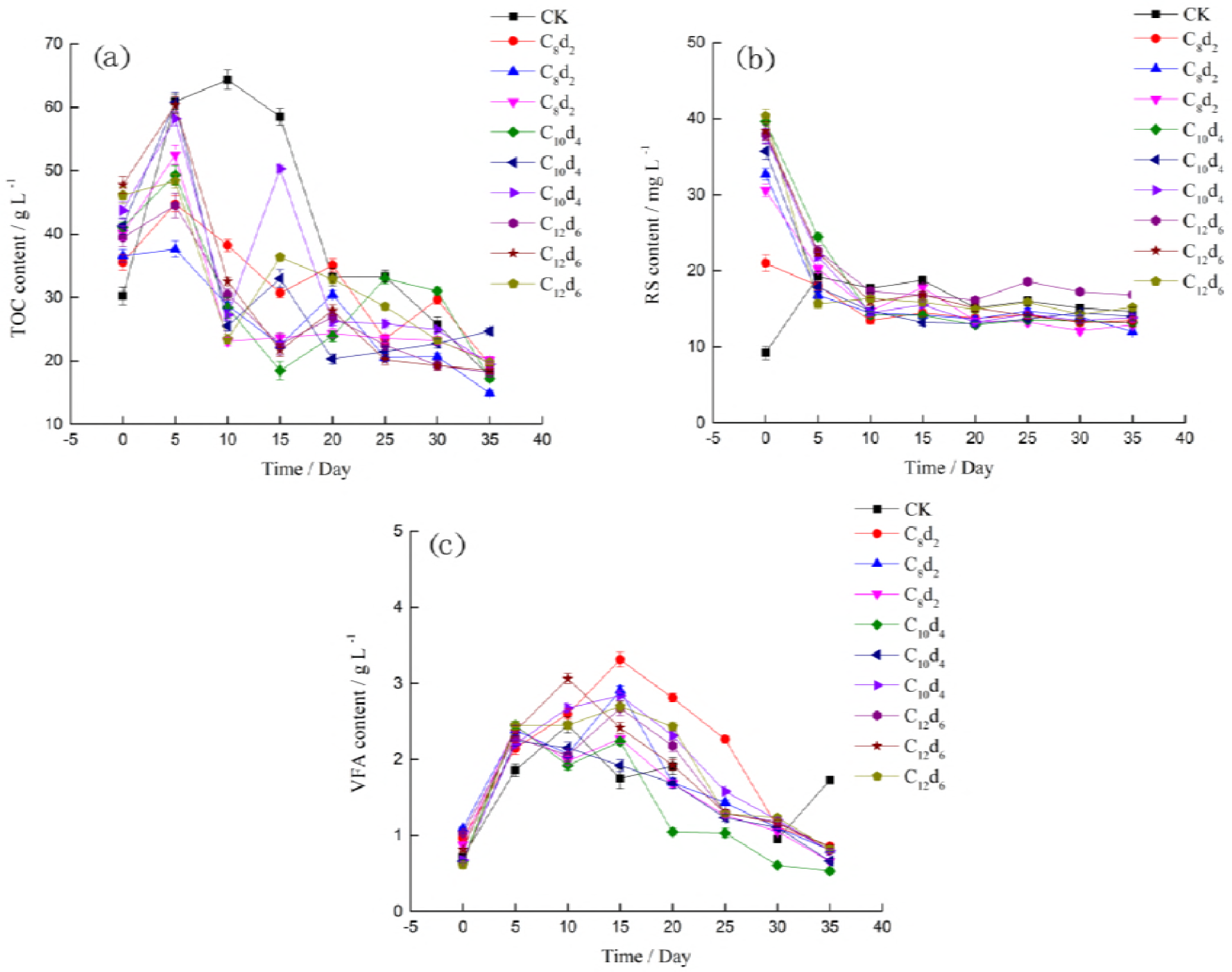
Nutrient substance content of AD process under NH_3_·H_2_O pretreatment. Note: CK is un-pretreated sample; C and d are the NH_3_·H_2_O pretreatment concentration and NH_3_·H_2_O pretreatment time. VFA means volatile fatty acid.

The figure 5 (b) indicated that the highest reducing sugar content of CK was 19.24 mg/L, and its built-up time appeared on the 5th day, while they were 21.01 mg/L, 32.73 mg/L, 30.59 mg/, 39.61 mg/L, 35.75 mg/L, 37.75 mg/L, 37.93mg/L, 38.40mg/L, 40.35mg/L, in the samples of C_8_d_2_, C_8_d_4_, C_8_d_6_, C_10_d_2_, C_10_d_4_, C_10_d_6_, C_12_d_2_, C_12_d_4_, C_12_d_6,_ respectively, their appearance time appeared on days 0. The results indicated that the highest reducing sugar content of each pretreated sample was higher than that of CK in AD process, and its appearance day was enhanced. The probable reason may be the hydrolysis produced reducing sugar [41], at the same time the reducing sugar was consumed by microorganism in AD process [42], which indicated that the reducing sugar content in AD process was depended on the reducing sugar content of production minus consumption, which was stronger in NH_3_·H_2_O pretreatment sample than that in CK. Thus, the highest reducing sugar contents of pretreated samples were higher than that of CK during AD process.

The highest VFA content of CK was 2.49 mg/L during AD process, and its built-up time appeared on the 25th day (figure 5c). The highest VFA contents of C_8_d_2_, C_8_d_4_, C_8_d_6_, C_10_d_2_, C_10_d_4_, C_10_d_6_, C_12_d_2_, C_12_d_4_, C_12_d_6_ were 3.31 mg/L, 2.91 mg/L, 2.43 mg/, 2.45 mg/L, 2.42 mg/L, 2.84 mg/L, 2.67 mg/L, 3.06g/L, 2.70mg/L, they appeared on days 15, 5, 5, 5, 5, 15, 15, 10, 15, respectively. Thus, the results concluded that the NH_3_·H_2_O had an effect on the first peak values of VFA content and their built-up time that was advanced. For the reducing sugar produced by NH_3_·H_2_O pretreatment could be decomposed into VFA by microorganism, immediately (Figure 4). While the reducing sugar during AD process of CK was produced by hydrolytic action and then followed by VFA[43], so the highest VFA content of pretreated sample was advanced during AD process (see figure 5).

The variation tendency of the contents of TOC, reducing sugar and VFA were affected by NH_3_·H_2_O. In CK, the variation tendency was that the TOC content was increased in the first 10 days, and it decreased in the days 10 to 20, after days 20, it decreased slowly. The reducing sugar content was increased in the first 5 days, and it decreased in the days 10. And there were two obvious peak values of the VFA content in AD process, after days 25, it decreased slowly. In liquid stage, the complex organic matter was decomposed into dissolved organic matter and then the dissolved organic mattes were metabolized to monosaccharide [43]. Thus, the content of TOC and reducing sugar was increased in the first 5 days. As time went on, the reducing sugar was decomposed to VFA in acid stage, the VFA content increased[44]. At last, the VFA was dissolved by methane bacteria to produce biogas, the VFA content decreased [35]. In the pretreated samples, the variations tendency of the TOC were increased to the peak value in 5 days then decreased irregularly, after 20 days the variation tendency of the TOC was small fluctuation. The variation tendency of the reducing sugar was decreased in the first 10 days, and after 10 days the variation tendency had a little change. Similarly, the VFA content of pretreated samples were decreased slowly after the days of their peak values. Thus, we concluded that variation tendency of the TOC content, reducing sugar content and cellulase activity had a big fluctuation in the earlier stage, maybe because the NH_3_·H_2_O pretreatment changed the initial digestion environment, which made the start time of hydrolysis stage advanced and the hydrolysis stage and acidification stage started, concurrently (figure 4). As time went on, the TOC and VFA was consumed by microorganism, the power of NH_3_·H_2_O pretreatment is reduced, and the differences decreased between CK and pretreated samples. Furthermore, the production rates of TOC, reducing sugar and VFA were the same as their consumption rates, the AD system had a dynamic balance. Thus, the content of TOC, reducing sugar and VFA had small fluctuation in last stage. The variation tendency of reducing sugar was smaller than that of TOC and VFA, may be because reducing sugar was quickly decomposed by microorganism into VFA and reducing sugar was not accumulated in AD process, thus, the variation tendency of reducing sugar was more stable.

#### 3.3.2 The influence of NH_3_·H_2_O pretreatment on the micro-ecological factor in AD process

As shown in figure 6 (a), the maximum cellulase activity of CK was 6.19 IU/mL, and its built-up time was the 10 days, while that of C_8_d_2_, C_8_d_4_, C_8_d_6_, C_10_d_2_, C_10_d_4_, C_10_d_6_, C_12_d_2_, C_12_d_4_, C_12_d_6_ were 5.54 IU/mL, 4.66 IU/mL 5.10 IU/mL 4.06 IU/mL, 4.38 IU/mL, 6.11 4.37 IU/mL, 5.94 IU/mL and 6.10IU/mL, respectively, each of then appeared on day 0. It was indicated the variation tendency of the cellulase activity was affected by NH_3_·H_2_O. In CK, the variation tendency was that the cellulase activity content was increased in the first 10 days, and it decreased to the lowest in 15 days then increased again. Because, there was a relationship between cellulase activity and stage of hydrolysis in AD process [45], the cellulase activity was increased in stage of hydrolysis to produce more soluble organic matter, but the rise of the soluble organic matter had a reaction on the cellulase activity, so than the cellulase activity decreased [46]. As the same principle, the NH_3_·H_2_O pretreated made the soluble organic matter and the cellulase activity decreased in the first 5 days, then the cellulase activity decreased. After 25 days the variations tendency of the cellulase activity in CK and pretreated samples were the same and had a small fluctuation. The reason was because the reducing sugar was quickly decomposed by microorganism into VFA and reducing sugar was not accumulated in AD process (figure 5).

**Figure 6.**
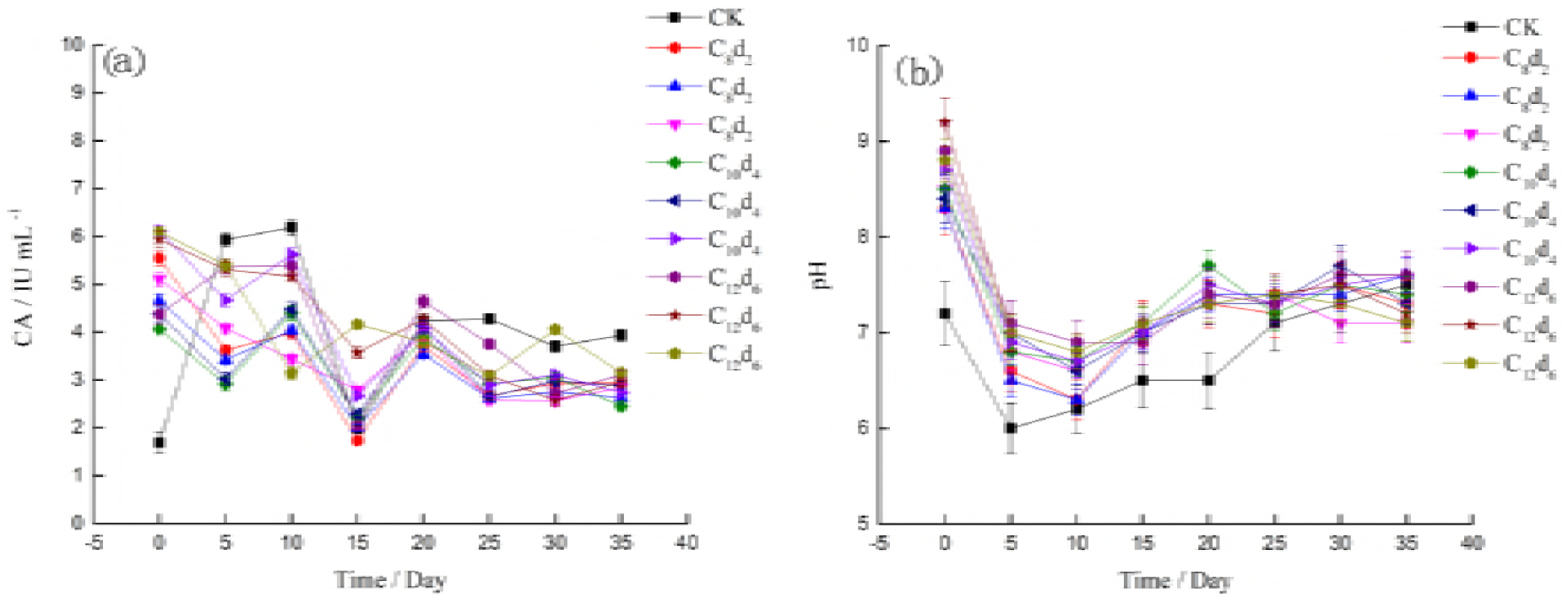
pH content of AD process under NH_3_·H_2_O pretreatment. Note: CK is un-pretreated sample; C and d are the NH_3_·H_2_O pretreatment concentration and NH_3_·H_2_O pretreatment time.

As shown in figure 6 (b), the maximum pH of CK was 7.5 and its built-up time appeared on the 35th day. The maximum pH of C_8_d_2_, C_8_d_4_, C_8_d_6_, C_10_d_2_, C_10_d_4_, C_10_d_6_, C_12_d_2_, C_12_d_4_, C_12_d_6_ were 8.3, 8,3, 8.5, 8.5, 8.4, 8.7, 8.9 and 8.8 respectively, they all appeared on days 0. The minimum pH of CK was days 10, the same with the days of pretreated samples in AD. It was indicated that all of the maximum pH of pretreated samples in AD process were higher than that of CK, and their appearance days was advanced as the NH_3_·H_2_O is an alkaline reagent. The changed trend of pH was not affected by NH_3_·H_2_O. In CK, the variation tendency was that the pH was decreased in the first 5 days, and it increased slowly. In the pretreated samples, the variations tendency of the pH was the same as CK. At the beginning of the AD, VFA content was low, then VFA was produced by hydrolysis and acid bacteria, thereby increasing the VFA content and decreasing the pH. Along with the digestion, VFA, acid, and dissolved nitrogen-containing compounds were broken into CH_4_, ammonia, amines, carbonates, and other molecules (CO_2_, N_2_, CH_4_, and H_2_), thereby decreasing the VFA content and increasing the pH [17].

#### 3.3.3 The influence of NH_3_·H_2_O pretreatment on the stability of environmental factors in AD process

In the table 2, the CVs of TOC content, reducing sugar content, VFA content, cellulase activity, pH, DBP in CK and pretreated samples were showed, they were affected by NH_3_·H_2_O pretreatment. The lower the stability, the greater the CVs were [47]. Thus, we concluded that the NH_3_·H_2_O pretreatment could make the stability of TOC, reducing sugar, VFA content cellulase activity and pH lower than that of CK. May be because their standard deviations were bigger than that of CK. However, the stability of DBP was increased by NH_3_·H_2_O pretreatment for the higher average value and lower standard deviations that was because activation speed of AD is accelerated and the time of larger DBP is longer, so the CVs of pretreated samples were reduced and stability rose. From the result that the stability of nutrient substances and environmental conditions were reduced while the stability of DBP increased, we concluded that ecological factors in AD system had negative effect on DBP. We thought may be because the nutrient substances and environmental conditions during AD process works with each other to make the digestion environment more suitable for AD [48], but this point need we do a further analysis.

**Table 2.**
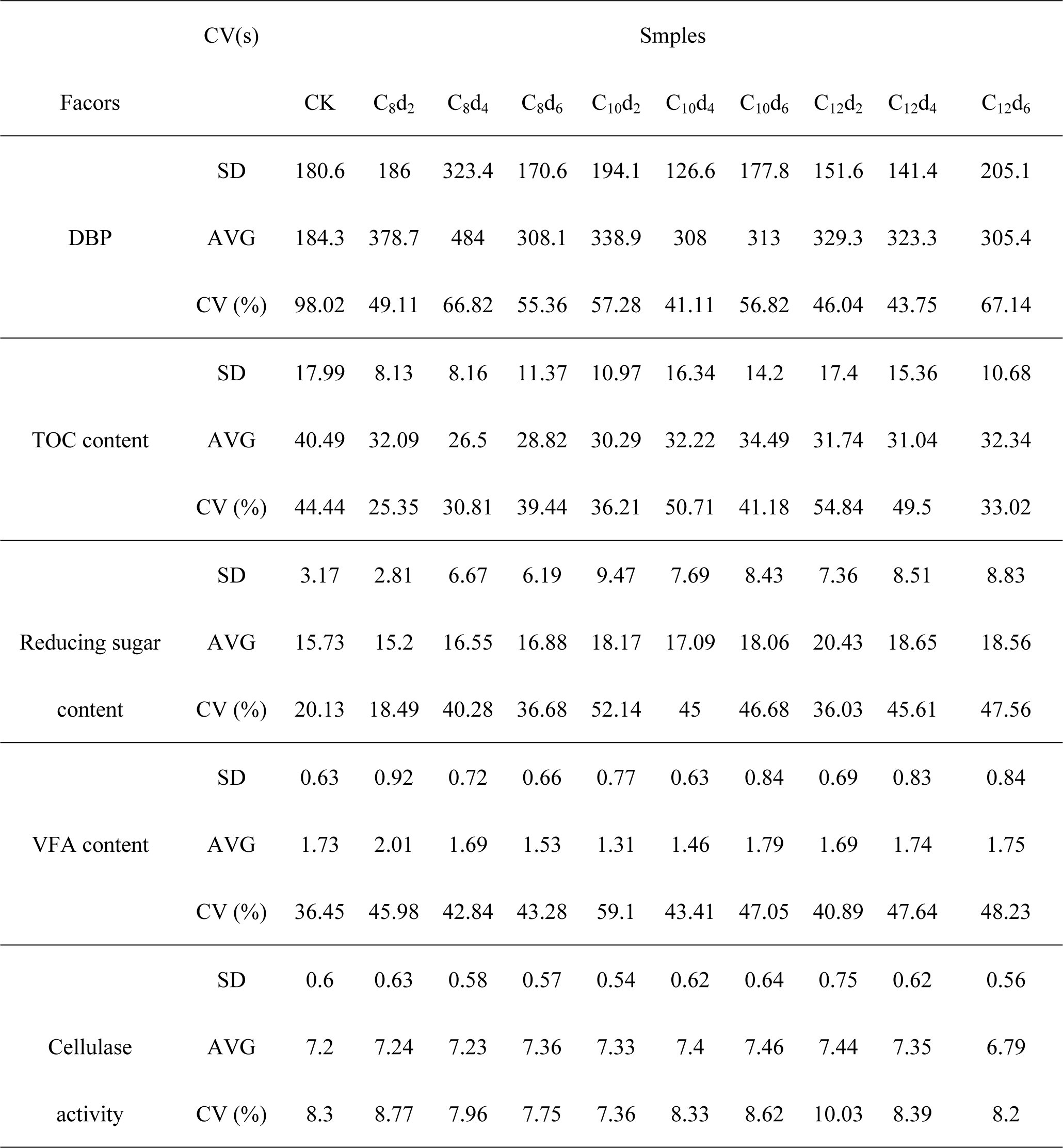
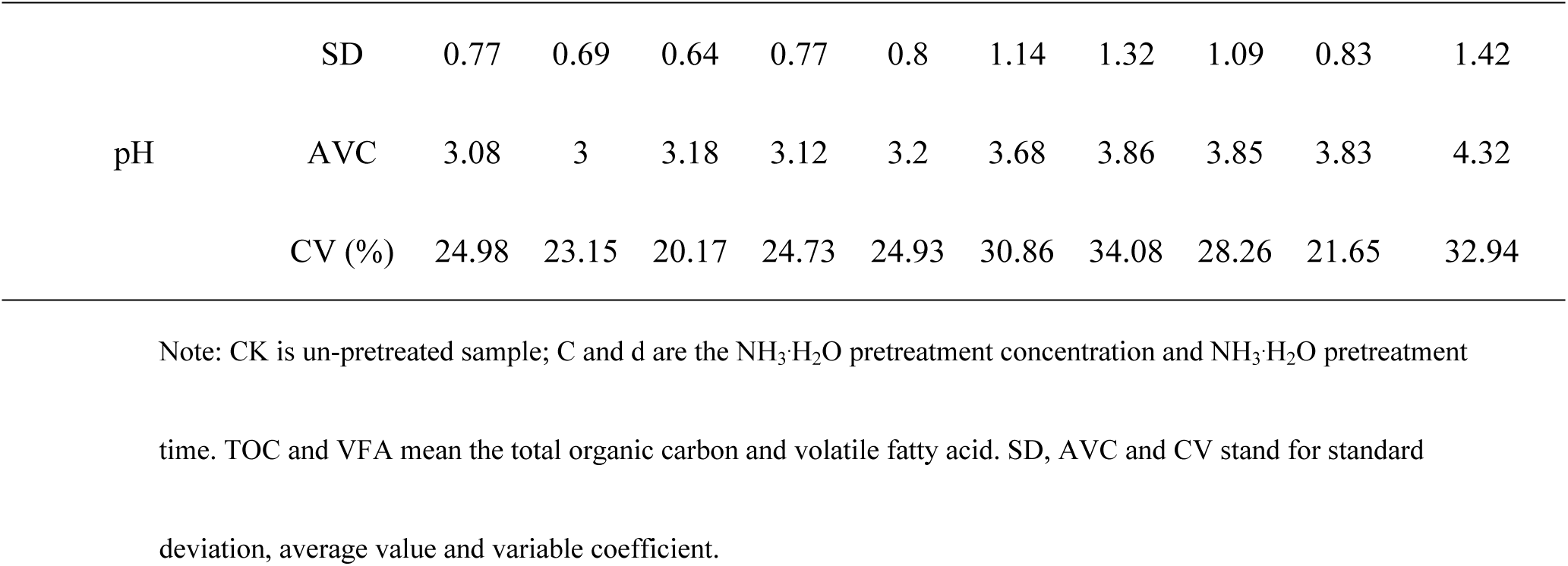
the CV of environmental factors in AD process under NH_3_·H_2_O pretreatment

### 3.4 The effect of the environment factors in AD on the biogas production

#### 3.4.1 The effect of environment factors on the stability of DBP

In statistics, stability was expressed by CV [27], thus the effect of environment factors on the CV of the DBP could indicated the effect of environment factors on the stability of DBP during AD system. The effect of the initial AD liquid characteristics on the CVs of environment factors during AD system were analyzed by RDA and showed in figure 7. Monte Carlo analogy was used in RDA by 999 times. The P value was 0.039 that was less than 0.05, thus there was a significant correlation between the model and the real value with statistical significance. The result showed that the first axis could explain 88. 5% of all information and second axis could explain 6.45%, thus the total rate explained from first axis and second axis was 85.83%, which indicated that the first axis and second axis could perfectly reflect the relationship between the initial AD liquid characteristics and CVs of environment factors during AD system, perfectly, and the first axis had the most decisive role. The cellulase activity had the biggest negative influence on the CV of TOC content pH, while it had the biggest positive influence on the CV of reducing sugar content and VFA content. As well as the VFA content had negative influence on the CV of reducing sugar content and TOC content, while it had the most positive effect on the CV of pH. Additional, the reducing sugar content, pH, TOC content and VFA content had positive effect (pH had the biggest) on CV of cellulase activity, while they had the negative effect on the CV of DBP. Thus, we concluded that the initial environment factors had influences on the stability of environmental factors. Because the environment factors characteristics (maximum value, minimum value and their appearance time) during AD system were affected by initial environment factors including substrate characteristics [22], pretreatment modes [21], initial liquid environment characteristics [3] that effected the stability of AD system. Although, the initial environment factors had positive effect or negative effect on the stability of environment factors in AD system, initial environment factors (the TOC content, RS content, VFA content, cellulase activity and pH) had the positive effect on the stability of DBP during AD process. The possible reason may be that the initial environment factors made the environment factors interaction with each other, which enhance the stability of system, so the stability of DBP was enhanced. Thus, response relationship between the stability of environmental factors and TBP was needed further study.

**Figure 7.**
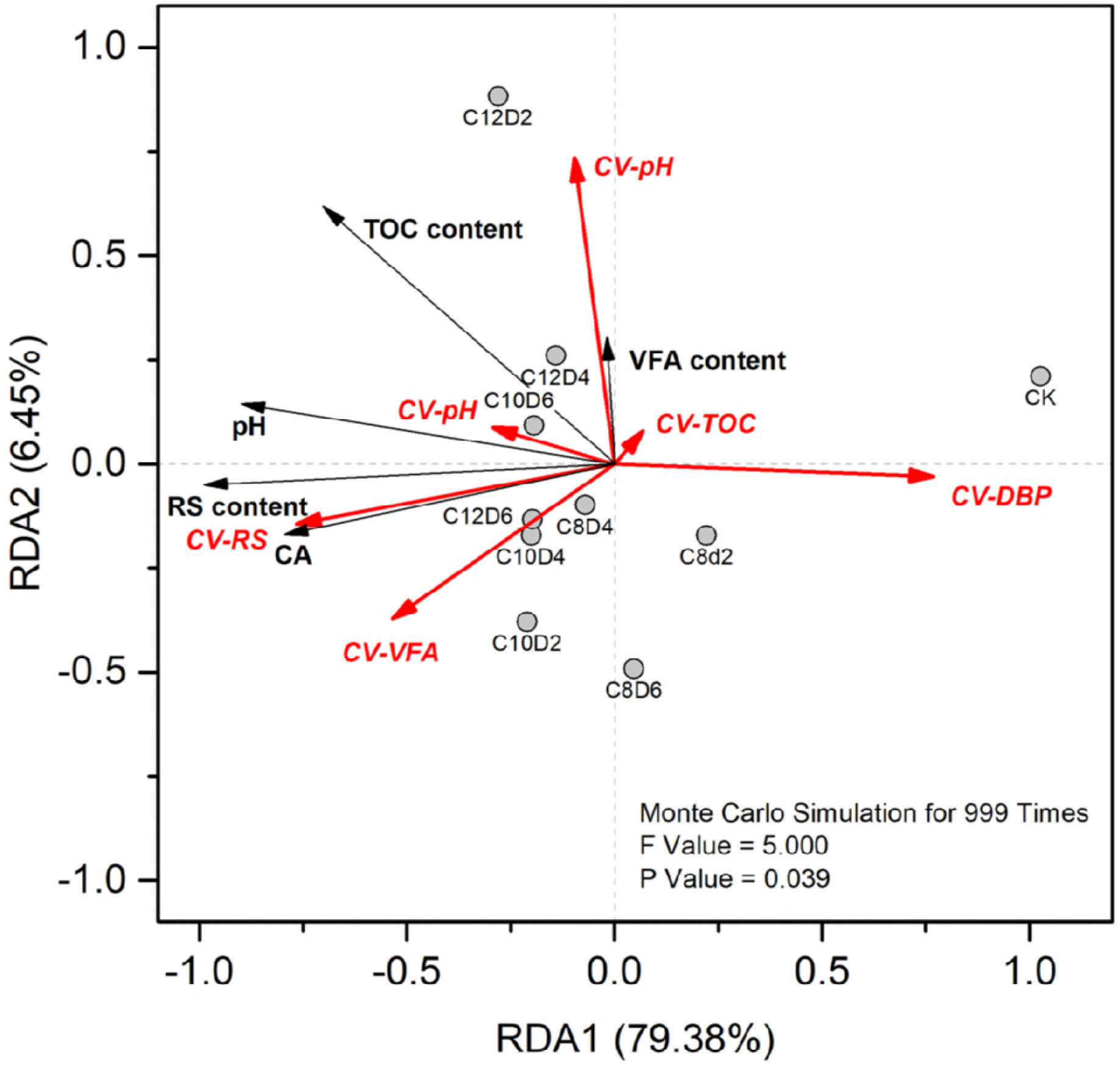
Effect of initial environment factors on the CV of environmental factors. Note: CK is un-pretreated sample; C and d are the NH_3_·H_2_O pretreatment concentration and NH_3_·H_2_O pretreatment time. TOC and VFA mean the total organic carbon and volatile fatty acid. CV stands for variable coefficient.

#### 3.4.2 The effect of the system stability of AD system on TBP

Path analysis was used to examine the direct and indirect influences that could reflect the relationships between the CV of environmental factors in AD process (TOC content, reducing sugar content, VFA content, cellulase activity and pH) and TBP, the data was showed in table 6 that indicated that the CV of environmental factors in AD process (TOC content, reducing sugar content, VFA content, cellulase activity and pH) had direct and indirect influences on the TBP, because changes of the CV of TOC content, reducing sugar content, VFA content, cellulase activity and pH correlation during AD process made the environment factors coherent on with another, which made the digestion environment more suitable for microbial life and more successful. And the CV of reducing sugar content had the highest direct influence on TBP, while the CV of TOC content had the highest total influence on TBP. The probable reason may be that the reducing sugar was the common product of intermediate metabolism of many kinds of microorganisms[49] and it also was the direct matter of decomposition of cellulose by many kinds of microorganisms[50], thus the CV of reducing sugar content had the highest direct influence on TBP, while the TOC was decompose into reducing sugar that was decompose into VFA, following, and the VFA had effect on the pH during AD process, which concluded that the TOC could effected the TBP by the environment factors, such as reducing sugar, VFA and pH, so the CV of TOC had the highest total effect on the TBP. The CVs of environment factors during AD process reflected the stability of AD system. Thus, the path analysis results indicated that stability of AD system affected the TBP. Previous study has shown that pretreatment improve the stability of AD system to enhance the TBP [51], but its measurements were the suitable ranges of environment factors during AD process, such as VFA content, alkalinity, ammonium nitrogen content, which were used by previous study to explain the relationship between stability of AD system and TBP. Moreover, Yin [52] showed that the environment factors during AD process had direct and indirect influences on the TBP, and the pH had the most effect on TBP, which different from this result. Because the pH was the values that appeared in the AD process, while they were the CVs of pH in this study. The remaining path coefficient was 0.717 which indicated that other factors also affected TBP and the CVs of other environmental factors had the highest total influences on TBP. For example, alkalinity [53], ammonium nitrogen [54], C/N [55], ionic potential [56], chemical oxygen demand [57] and so on. The remaining path coefficient was 0.717 that was main the CV of environment factors during AD process had small effect on the TBP, because the environment factors had effect the TBP by co-operate fully with each other.

**Table 3.**
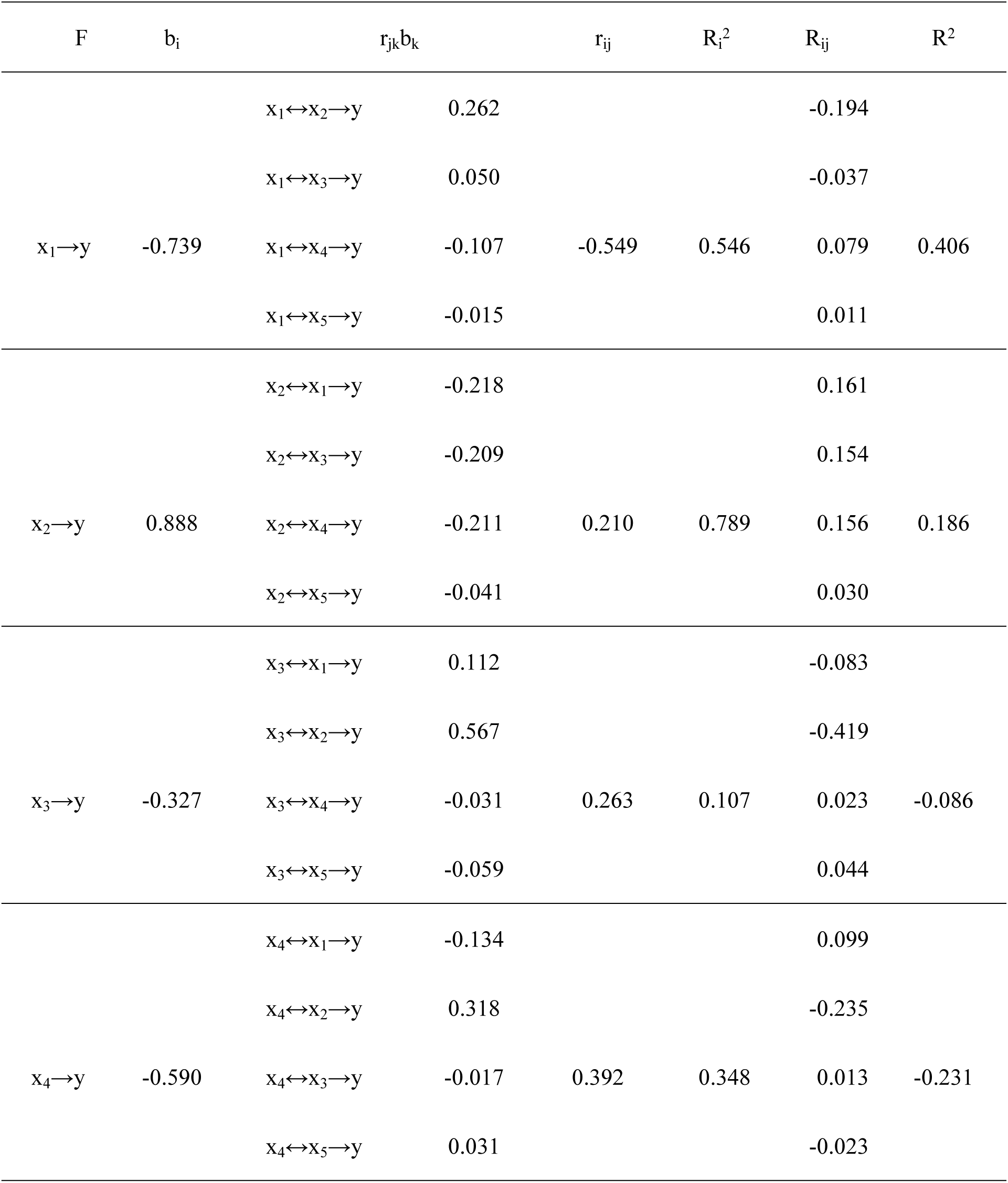
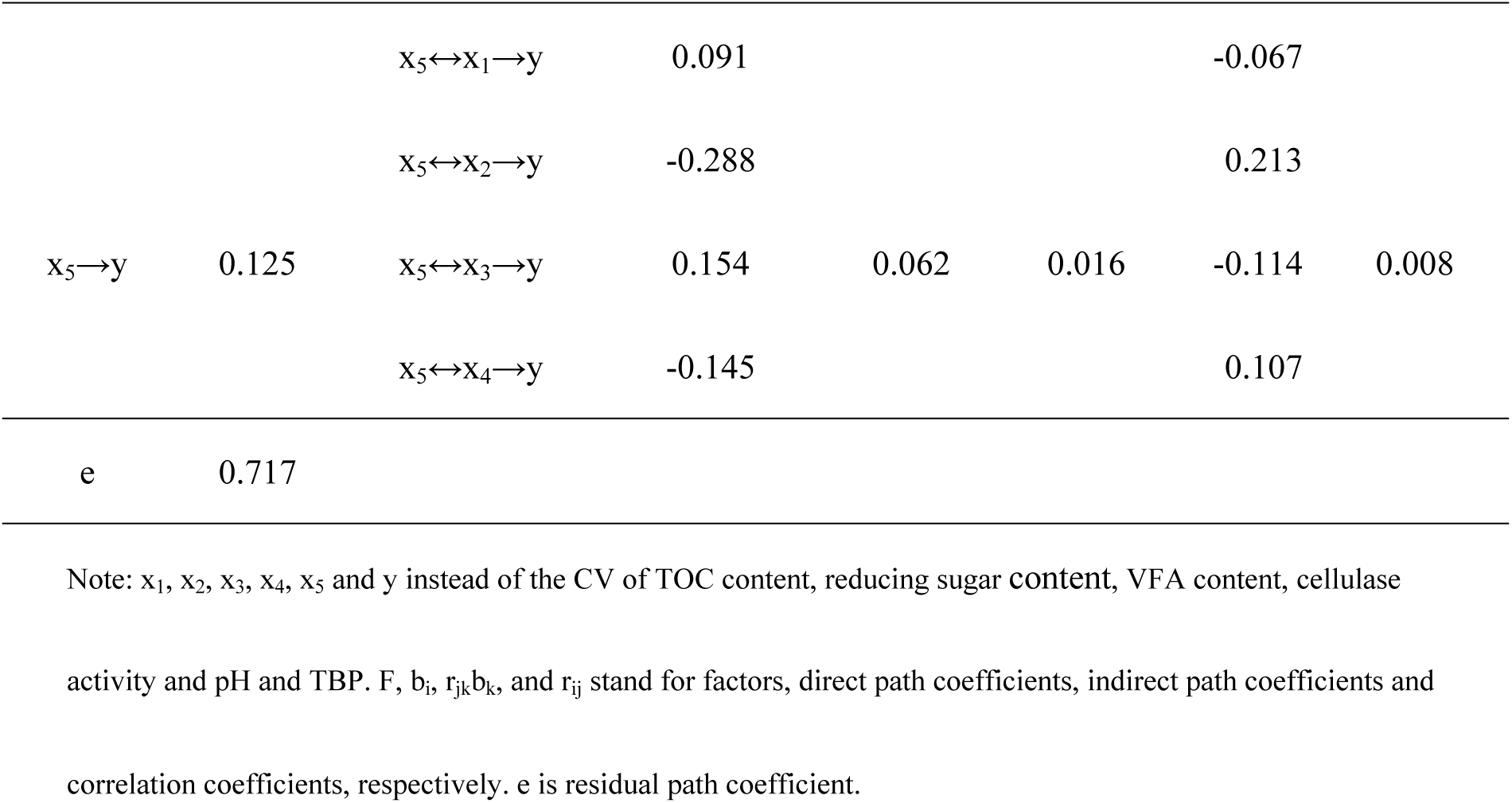
the response relationship between the CV of environmental factors and TBP

### 3.5 Reasons for improvement of TBP

The result showed that TBP was enhanced in AD after NH_3_·H_2_O pretreatment (See figure 3). Five possible reasons explained why NH_3_·H_2_O pretreatment improved TBP. First, the contents of TOC content, reducing sugar content and VFA content in NH_3_·H_2_O pretreated samples were higher than that of CK, TOC, reducing sugar and VFA was the nutrient substance that can increase microbial activity and bring forward the start-up time of the AD system to make the highest DBP improved [35]. Second, the cellulose activity was higher than in CK, and the decomposition of cellulose was accelerated (figure 3), which could deliver nutrients quickly to adjust the growth of methane bacteria when the cellulose activity increased [59]. Third, NH_3_·H_2_O pretreatment increased the pH of the initial digestion liquid and advanced the acid phase (figure 3) [58]. Fourth, the stability of DBP was enhanced, as well as the appearance time of the highest DBP was appeared advanced and the average value of DBP was increased by NH_3_·H_2_O pretreatment. The last remaining factors in the initial digestion environment affected the environmental factors that affected the AD stability of AD system, and the environmental factors had direct and indirect influences on the TBP, which can coordinate environmental factors in AD system to be more suitable for AD and the AD system can be more stability.

## 4 Conclusions

Pretreatment at 8% for 4 days, the TBP produced the highest value (302.5mL/g TS), which was 263% higher than that of CK (115.18 mL/gTS) and significantly higher than the other values (P < 0.01). NH_3_·H_2_O pretreatment had effect on the initial AD environment factors and the environment factors during AD process. Under the NH_3_·H_2_O pretreatment conditions, the stability of environment factors during AD process was affected by initial AD environment factors, and the stability of environment factors during AD process had direct and indirect influences on the TBP. The main reason that why NH_3_·H_2_O pretreatment could improve the TBP was that NH_3_·H_2_O pretreatment affected the stability of environment factors during AD process by affecting the initial AD environment factors, as well as the stability of environment factors had a cooperation action among each other, which made the environment during digestion process more suitable for AD and improved the stability of AD system.

## Acknowledgments

This work was supported by Sichuan Provincial Science and Technology Department Application Foundation Fund (2014JY0196).

**Abbreviations**
NH_3_·H_2_O: Ammonium hydroxide
TBP: Total biogas production
AD: Anaerobic digestion
RS: Rice straw
DM: Dairy manure
TS: Total solid
VFA: Volatile fatty acid
DBP: Daily biogas production
TOC: Total organic carbon

